# EXPERIMENTAL INVESTIGATION OF PULSE STERILIZATION OF VIRAL INFECTION

**DOI:** 10.1101/2020.12.16.423002

**Authors:** Volodymyr Chumakov, Mykhailo Ostryzhnyi, Oksana Kharchenko, Krystina Naumenko, Svitlana Zagorodnya, Vasiliy Muraveinyk, Aleksandr Tarasevich

**Affiliations:** Kharkiv National University of Radioelectronics, Kharkiv, Ukraine; Danylo Zabolotny Institute of Microbiology and Virology of the National Academy of Science of Ukraine, Kyiv, Ukraine; limited Liability Company «Triix», Chernihiv, Ukraine

## Abstract

The results of experimental investigations of the effect of high-intensity pulsed UV radiation on the influenza virus type A (H1N1) are presented. The research methodology is developed and the structure of the experiments is described. An end-face plasma accelerator was used as a radiation source, which provides a power pulsed discharge in an open atmosphere. The high efficiency of inactivation of the infectiousness of the virus was shown within a short period of time. The possibility of providing urgent 100% sterilization of a viral infection has been shown for the first time. A model for calculating the efficiency of pulse sterilization has been developed. The prospects for the application of pulse sterilization technology to combat coronavirus infection are considered.

## Introduction

The large-scale pandemic caused by the SARS-CoV-2 virus has largely updated research in the development and creation of methods and means of anti-epidemic protection of the population. Along with pharmacological agents and vaccination, physical methods of antiviral antibacterial treatment of the human environment, based on the use of ultraviolet (UV) radiation, open up broad prospects [1–4].

The vast majority of UV sources of continuous radiation flux have a number of fundamental disadvantages due to their low power. So, for example, the power flux density created by the most widespread mercury germicidal lamps is of the order of milliwatts per square centimeter [5]. This leads to a significant increase in the duration of the process of antiviral and / or antibacterial processing, which is caused by the need to accumulate a bactericidal dose of UV radiation required to achieve a given level of sterilization. And, what is more dangerous, prolonged exposure to low intensity is to a certain extent mutagenic, which leads to the formation of a clinical infection in the case of bactericidal microflora, or to the preservation of residual virulence of the treated infectious environment [6,7].

In such conditions, the advantages of pulsed UV radiation of high and ultra-high power levels are manifested, providing the effect of complete destruction of bactericidal microflora and viruses. For the first time, the effect of 100% sterilization was considered in [8–10].

This paper presents the results of an experimental study of the antiviral efficiency of a pulsed source of broadband UV radiation based on an open discharge in the atmosphere at the exit of an end-face plasma accelerator.

### 1. Preparing the experiment

#### 1.1. Cultivation and preparation of cells

Cells were grown in sterile plastic flasks (Sarstedt, Germany) in a nutrient medium consisting of 45% DMEM (Sigma, USA), 45% RPMI 1640 (Sigma, USA), and 10% fetal calf serum (FBS) (Sigma, USA), inactivated by heating for 30 min at T = 56°C and the antibiotic gentamicin (100 μg/ml). The cells were passaged until a monolayer was formed, namely, they were disintegrated from the surface of the vials using 0.02% Versen solution (BioTestMed, Ukraine) and 0.025% trypsin solution (BioTestMed, Ukraine), resuspended in a nutrient medium, after which their concentration in the suspension was brought to (2-2.5) ×10^5^ cells/ml. The seeding ratio was established after counting the number of cells in the Goryaev chamber using an inverting microscope (Carl Zeiss Jena, Germany) with a magnification of 70x. After that, a cell suspension in a volume of 200 μl was introduced into the wells of a 96-well plate of the scanner, and then the plates with cells were cultured in a thermostat at 37°C in an atmosphere of 5% CO_2_. After 24 h of cultivation, the state of the cell monolayer in the plates was monitored using a microscope. For the research, we used samples in which the formation of cells of about 90% of the monolayer was observed in the absence of bacterial and fungal contamination. In all other cases, cells were excluded from the study. A mixture of 50% DMEM (Sigma, USA) and 50% RPMI 1640 (Sigma, USA) was used as a supporting medium for the cells.

#### 1.2. Preparation of virus

The studies used influenza virus type A (H1N1) strain A / FM / 1/47, obtained from the collection of the State Institution of the Institute of Epidemiology and Infectious Diseases named after L.V. Gromashevsky NAMS of Ukraine. The virus was cultured in an MDCK cell culture and stored in aliquots at −70°C.

A monolayer of cells in 650 ml vials for 24 hours of growth was washed with sterile phosphate-buffered saline (pH = 7.4), after which the virus was introduced into the cell medium in an amount sufficient to cover the monolayer at a dilution of 1:10000. Then, after adsorption for 1.5 h at room temperature, the required amount of a supporting medium and cells with viruses were added, incubated at 37°C for 4 - 5 days until an intense cytopathic effect (CPE) appeared. To isolate the virus from the cellular material, 3-fold freezing-thawing was carried out until the cells were completely destroyed, and the cell detritus was removed by centrifugation at 3000 rpm for 30 min. Finally, the titers of the virus in the supernatants were determined using the MTT method; the blanks were poured into cryovials of 1–5 ml and stored at T = −70°C. In the study, a virus pool with a titer of 5×10^6^ TCID50/ml was used.

#### 1.3 Pulse radiation source

An end-type plasma accelerator was used as the radiation source, which forms a plasma-dynamic discharge in an open atmosphere. As is known, in such sources with a high-current discharge, the current exceeds 10^5^ A and the brightness temperature is about 15000 K [11]. The large energy input in the mode of microsecond pulses provides the value of the plasma flow enthalpy, which is much higher than the values achieved in other sources. The radiation power in the bactericidal band of the spectrum was about 1-10 MW, which makes it possible to implement the radiation dose necessary for the complete destruction of the infection during one pulse at a distance of tens of centimeters from the emitter [8–10].

The brightness temperature of plasmodynamic radiation sources is proportional to the square of the current; therefore, in the design of the radiator, a busless circuit was used to minimize parasitic inductance [12,13]. The block diagram of the installation is shown in Fig. 1.

**Fig. 1.**
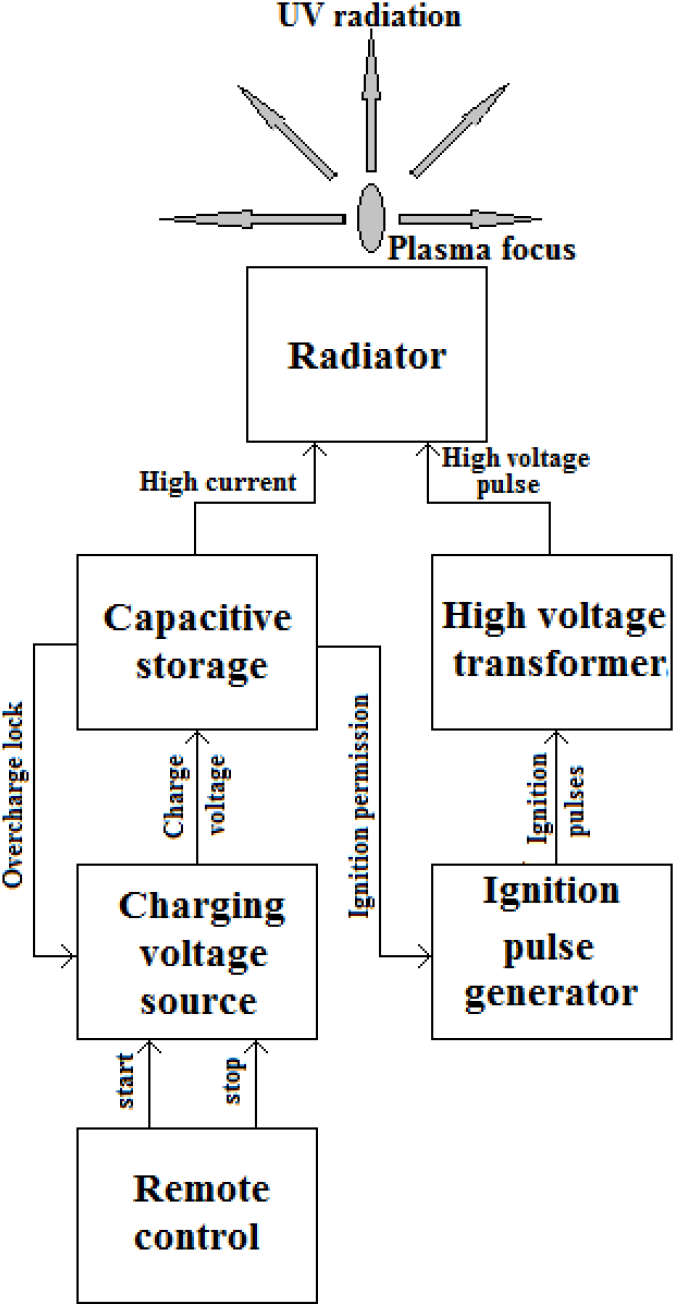
Structural scheme of the pulse UV radiation source

In fig. 2a shows a photograph of the discharge taken through an optical ultraviolet filter with a transmission band of 300 - 400 nm. As a result of image processing, the temperature distribution of the emitting region can be obtained (Fig. 2b) and the directivity characteristics of the radiation source can be estimated, similarly to [14].

**Fig. 2.**
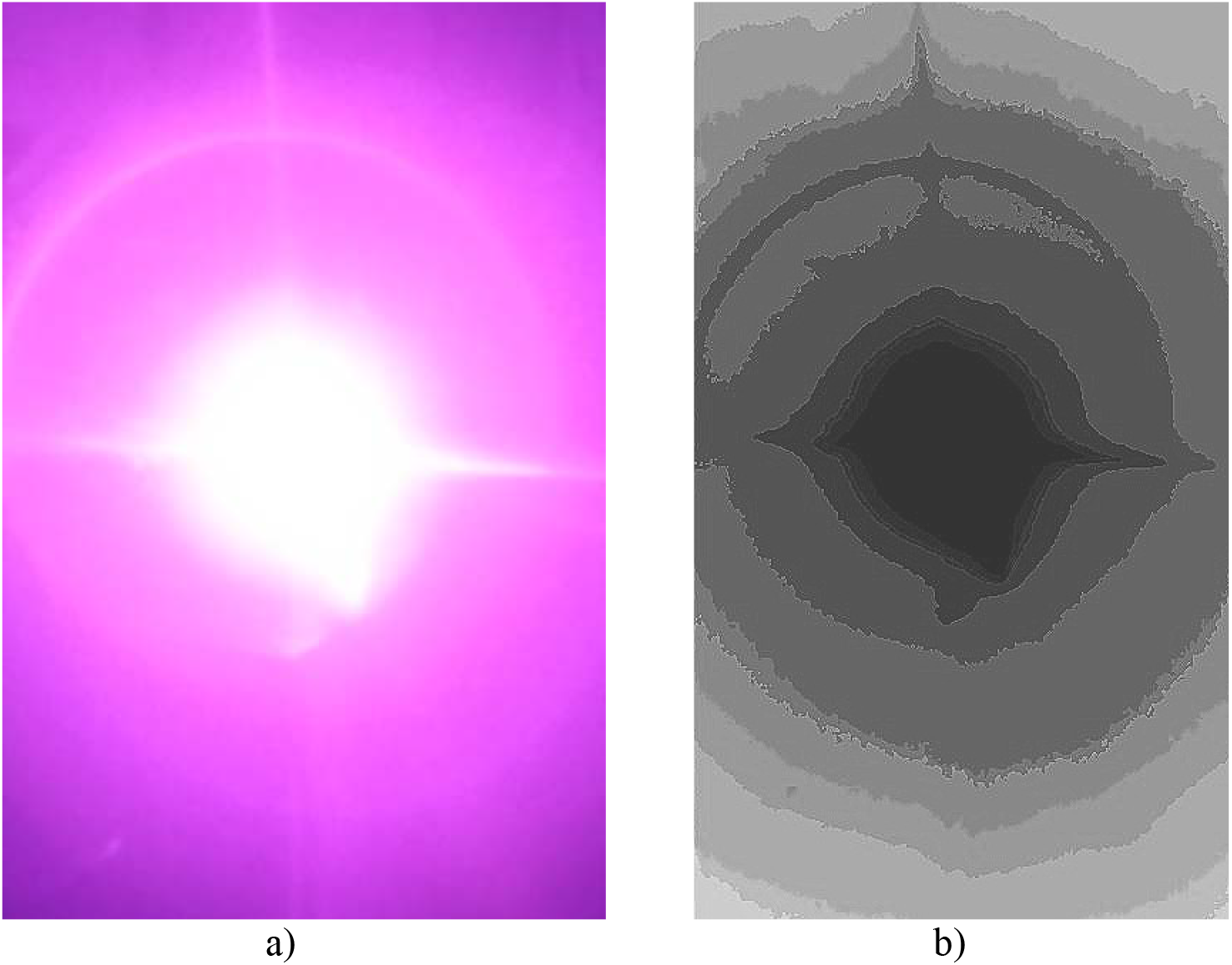
Plasma-dynamic discharge in the atmosphere (a) and temperature distribution (b)

### 2. Experiment and Results

The virus suspension in the cultivation medium was introduced into dishes, which were divided into 4 groups, and in the open state were placed in the irradiation zone of a pulsed UV radiation source (Fig. 3), after which they were irradiated. The radiation pattern of the emitter is close to uniform in the hemisphere, so each cup received the same radiation dose per pulse. Each of the groups of cups was irradiated with 1, 5, 10, and 16 UV pulses, respectively. Control sample - control virus (CV) was not exposed to radiation. Subsequently, the titers of infectivity of the control and experimental (exposed to irradiation) samples were investigated. For this, a 1-day 100% monolayer of sensitive cells was grown in a 96-well plate, which were infected with tenfold serial dilutions of the control and irradiated virus samples, 50 μl per well. The adsorption of the virus was carried out at a temperature of 37°C for 2 hours, after which 150 μl of the supporting medium was added to the wells. Cells not infected with the virus were used as control cells. Plates were incubated for 3 days in an atmosphere of 5% CO_2_ at 37°C until the appearance of pronounced CPE of the virus in the control sample.

**Fig. 3.**
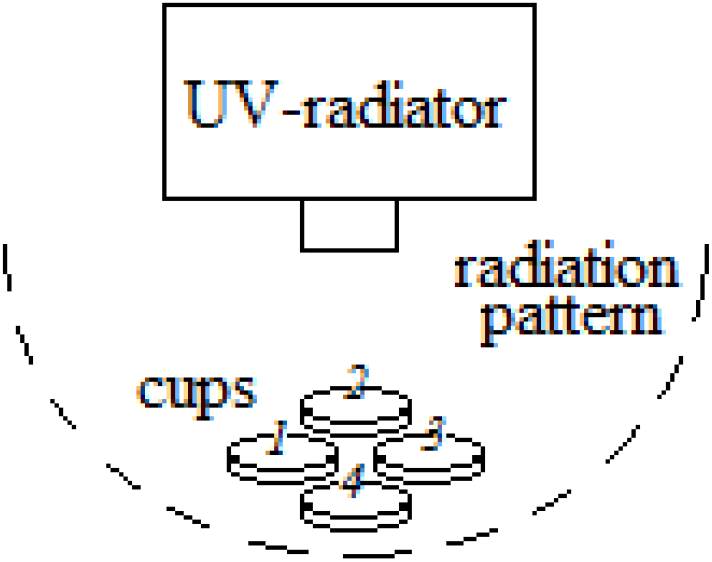
Experiment scheme

Crystal violet C25N3H30Cl was used for coloring. The absorption coefficient was measured using a Multiskan FC flatbed spectrophotometer (ThermoScientific, USA) at a wavelength of 538 nm. Using the measured values of optical density, the percentage of inhibition of cell viability under the influence of the virus (or % CPE of the virus on cells) was determined by the formula:

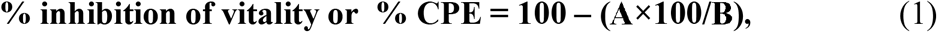

where A is the average optical density of the sample, and B is the average optical density of the control cells.

After that, the dilution of the virus was determined, which reduces the optical density of the sample under study compared to the optical density of the control cells by 50%, which is the titer of the virus and is expressed in TCID50/50 μl. Multiplying the obtained value by 20, the titer of the virus in 1 ml - TCID50/ml was determined, and then, expressing the obtained value in logarithms, the generally accepted expression of the titer of the virus logTCID50/ml was calculated.

The calculation of the index of suppression of the virus reproduction under the influence of radiation was carried out according to the formula:

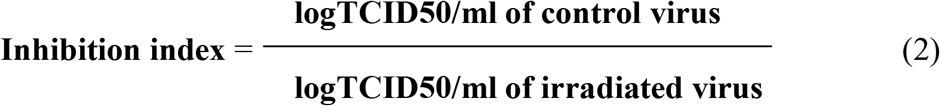

According to the effectiveness criteria, a decrease in the virus titer less than 1.25 is considered a relative effect, from 1.5 to 2.0 - a moderate effect, and more than 2.0 - a pronounced antiviral effect [15]. The results of the measurements and calculations are presented in table 1.

**Table 1 -.**
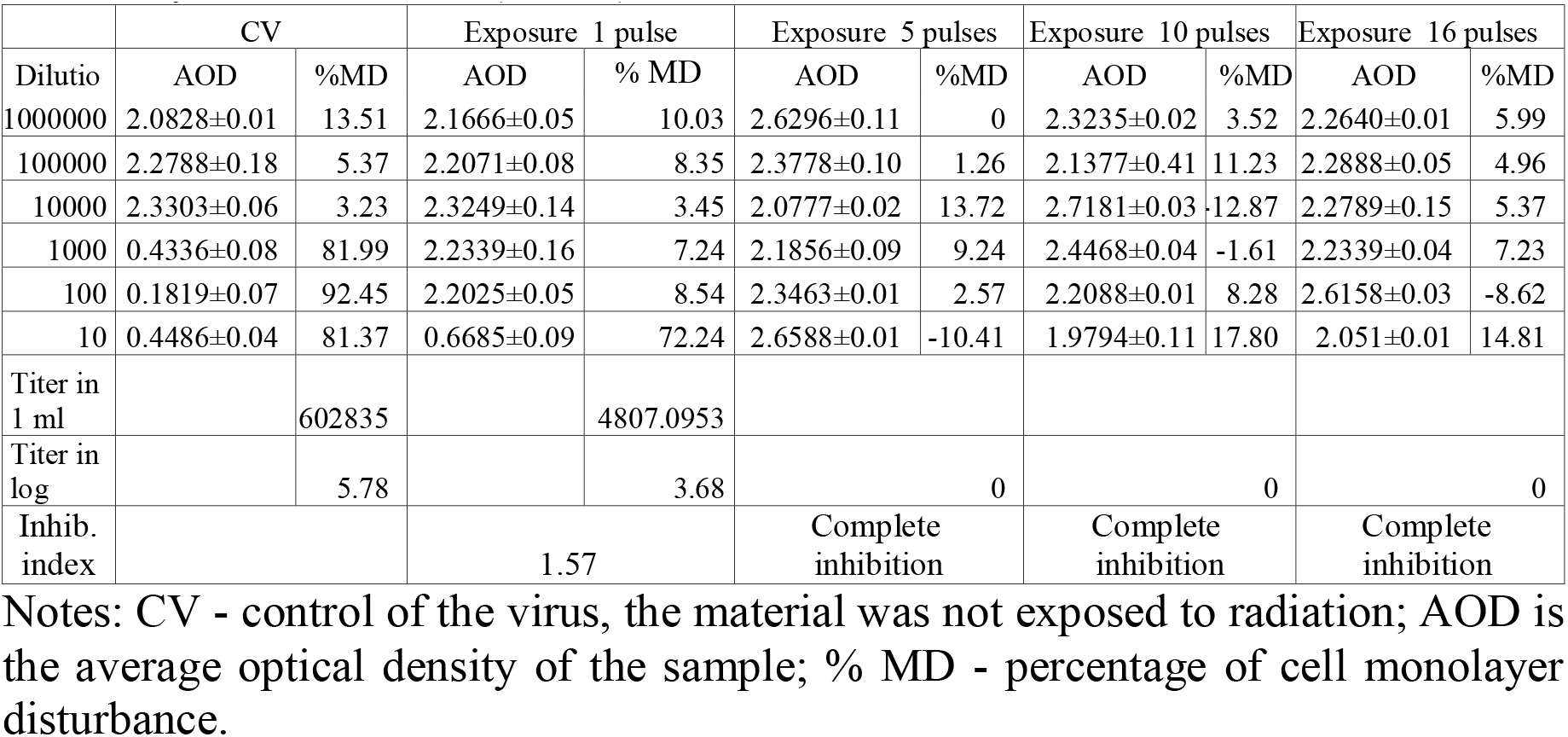
Results of the analysis of the effect of pulsed UV radiation on the infectivity of influenza A (H1N1) virus

The experimental dependence of the H1N1 virus inhibition index on the number of pulsed exposures is shown in Fig. 4.

**Fig. 4.**
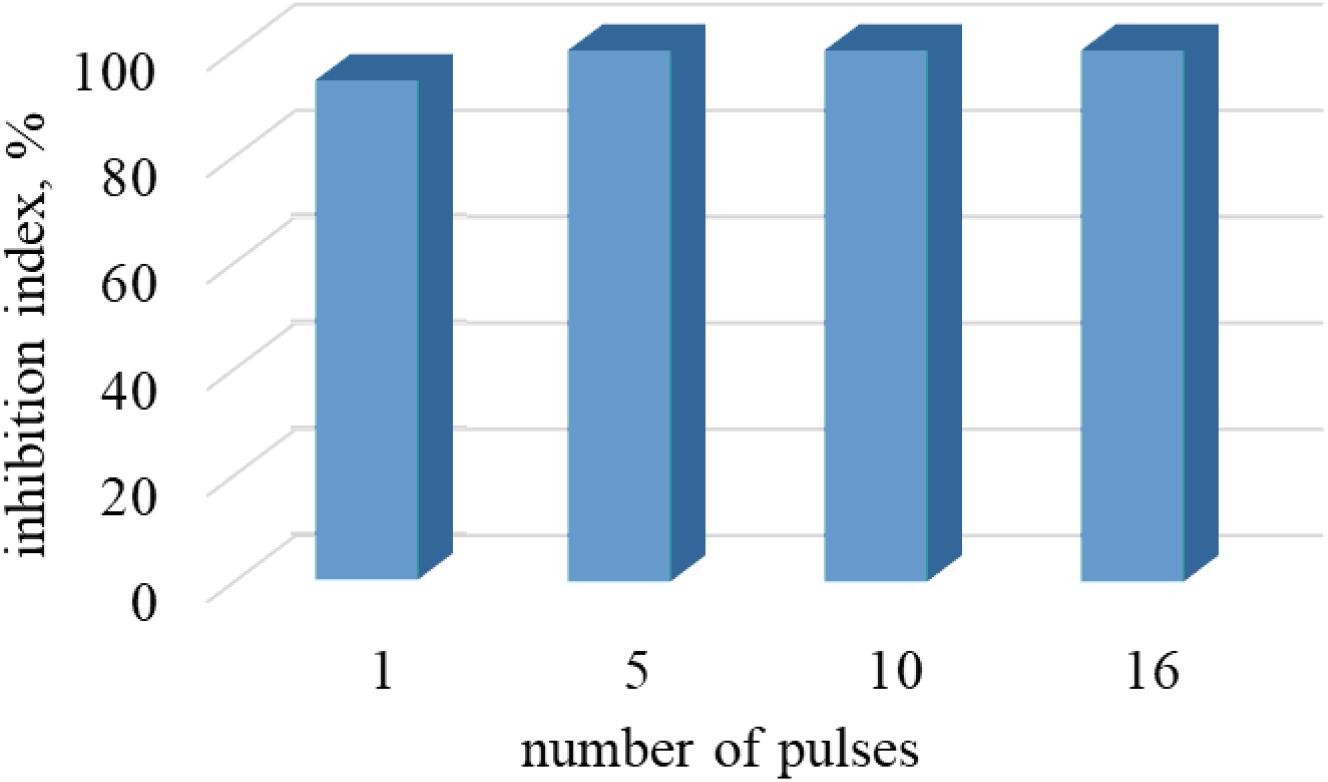
Oppression index dependence on number of pulses

As can be seen from the results of the study, as a result of irradiation with 1 pulse, the titer of the influenza virus significantly decreased (inhibition index 1.57), which corresponds to a sterilization efficiency of about 94%. After exposure to 5 or more pulses, the virus completely lost its infectivity, that is, the effect of complete sterilization is provided.

## 4. Discussion

When simulating the effect of continuous UV radiation, it is also assumed that the photobiological reaction of microorganisms and viruses is cumulative, as a result of which the bactericidal effect is determined by the radiation dose. Analytical models describing the dependence of bactericidal efficacy on the radiation dose are asymptotic in nature [16]. Sterilization efficiency, which is equal to the ratio of the number of microorganisms *N_s_* that survived after irradiation during exposure *t*, to their initial number *N*_0_ may be calculated

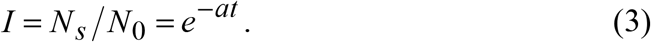

Formula (3) is usually written as

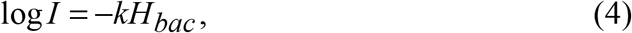

to show the dependence of bactericidal efficacy on the dose of radiation. The parameter *k* [m^2^/J] included in (4) characterizes the rate of sterilization and is determined by the power of the radiation source and the resistance of the irradiated microorganisms [17]. As can be seen, formula (4) expresses the Bouguer-Weber-Fechner law, which establishes a connection between physical impact and the reaction of a biological object [18].

Sterilization is a threshold effect, i.e. complete destruction of pathogenic microflora and/or viral infection occurs as a result of exposure to radiation, the dose of which is equal to *H_th_*. Thus, as follows from (3), (4) in a continuous mode to achieve a dose *H_bac_* = *H_th_*, the exposure time is infinite *t* = ∞

In the pulse mode, an expression for describing the kinetics of bacterial and viral populations under UV irradiation is described by the formula

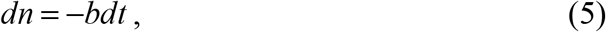

whence the bactericidal efficacy for the pulsed regime follows

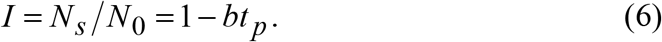

Here, the parameter *b* characterizes the duration of exposure at which complete destruction of the irradiated microorganisms takes place, and depends on the power of the radiation source, *t_p_* is the exposure time in the pulsed mode. Thus, according to [17], the parameter *k* in the model (4) determines the slope of the logarithmic curve of bactericidal efficiency.

Formulas (3), (4) and (6) make it possible to estimate the gain in processing time, which is provided in the pulse mode

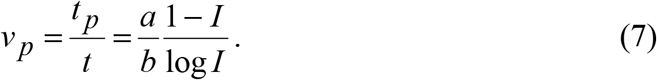

In fig. 5 a comparison of the exposure required for complete sterilization in continuous and pulsed modes is shown.

**Fig. 5.:**
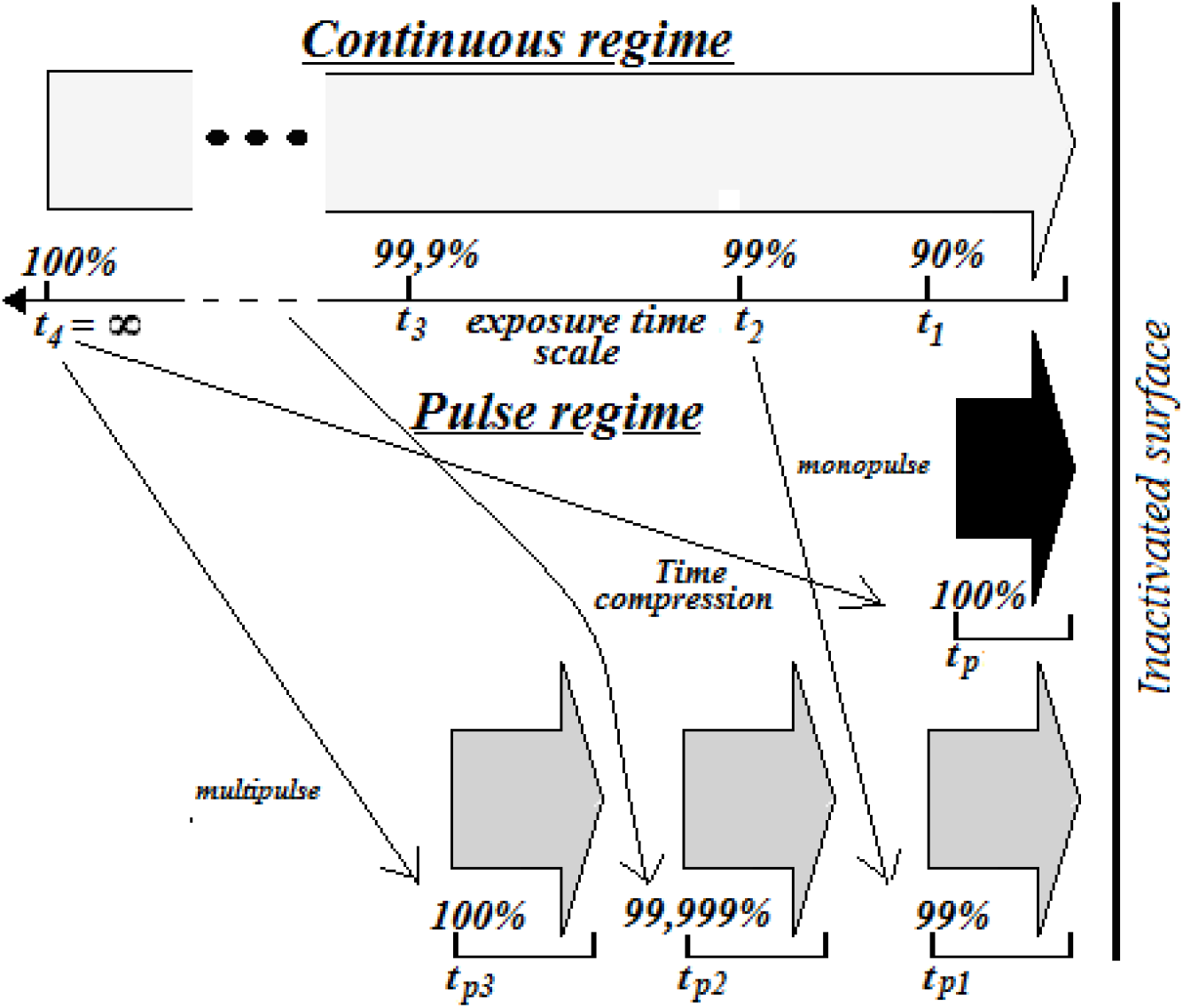
Comparison of exposure in continuous and pulsed modes

In the pulsed mode, the concentration of energy in a short time interval excludes the cumulative virucidal effect characteristic of continuous low-power exposures, and, more importantly, allows the required dose to be provided during one pulse. The action of a single pulse with a duration of *t_p_* = 30 μs provides a sterilization efficiency of about 94%. Complete sterilization is achieved with 5 pulses. Thus, the value of the gain (7) is *v_p_* ≈ 14222. In the calculations, we used *k* = 0.15 m^2^ /J, *H*_*bac*_0.94__ = 10 J/m^2^ [17.19], and the power of a continuous source *P_r_* = 30 W, *r* = 1 m.

Calculations and experimental measurements of the exposure time required to inactivate coronavirus infection seem to be very relevant today. The data on the resistance of viruses to UV radiation are very contradictory. For example, the known results of dose measurements of 99.9% inactivation of the SARS virus belonging to the same family as SARS-CoV-2 is 10–20 mJ/cm^2^ [20]. In [21], for the SARS-CoV-2 virus, the average dose of UV radiation, which provides a bactericidal efficiency of 90%, is *H_bac_* = 67 J/m^2^. On the other hand, in [22] the value of the threshold dose of UV radiation *H_bac_* = 36144 J/m^2^ is given, at which the destruction of SARS-CoV-2 with an efficiency of 99.999% is provided. Naturally, such significant discrepancies in the experimental data indicate the need for additional research. At the same time, such doses can be realized using one or several exposures of a pulsed radiation source used in this work.

## Conclusions

The results of the development and creation of a source of pulsed optical radiation based on a plasmodynamic system that forms a high-current discharge in an open atmosphere are presented. With such discharges, the power flux in the bactericidal band of UV radiation significantly exceeds the values that can be realized with the help of other sources of ultraviolet radiation.

Experimental studies of high-intensity pulsed UV radiation for influenza H1N1 virus have been carried out. Shown is the effect of reducing the level of infectivity of the virus during a short period of treatment. The basic principles of the mechanism of pulsed bactericidal and antiviral effects have been developed.

For the first time, the effect of complete inactivation of a viral infection under pulsed irradiation was obtained. The possibility of significant inhibition of viruses as a result of exposure to a single pulse of UV radiation with duration of 30 μs has been experimentally shown. The infection was completely inactivated by repeated pulsed irradiation. The results obtained open up broad prospects for the development of a pulsed technology for fighting infections, in particular, the development and creation of radiation sources that ensure the destruction of the SARS-CoV-2 virus.

